# Efficient discovery of potently neutralizing SARS-CoV-2 antibodies using LIBRA-seq with ligand blocking

**DOI:** 10.1101/2021.06.02.446813

**Authors:** Andrea R. Shiakolas, Nicole Johnson, Kevin J. Kramer, Naveenchandra Suryadevara, Daniel Wrapp, Sivakumar Periasamy, Kelsey A. Pilewski, Nagarajan Raju, Rachel Nargi, Rachel E. Sutton, Lauren Walker, Ian Setliff, James E. Crowe, Alexander Bukreyev, Robert H. Carnahan, Jason S. McLellan, Ivelin S. Georgiev

## Abstract

SARS-CoV-2 therapeutic antibody discovery efforts have met with notable success but have been associated with a generally inefficient process, requiring the production and characterization of exceptionally large numbers of candidates for the identification of a small set of leads. Here, we show that incorporating antibody–ligand blocking as part of LIBRA-seq, the high-throughput sequencing platform for antibody discovery, results in efficient identification of ultra-potent neutralizing antibodies against SARS-CoV-2. LIBRA-seq with ligand blocking is a general platform for functional antibody discovery targeting the disruption of antigen–ligand interactions.

---

Technologies for developing preventive and therapeutic measures that can counteract potential pandemics are of utmost significance for public health. The ongoing COVID-19 pandemic, for example, has emphasized the importance of developing effective countermeasures quickly. Through various pandemic preparedness initiatives, effective SARS-CoV-2-neutralizing antibodies were discovered and validated within months^1-7^, as were SARS-CoV-2 vaccine candidates^8^. However, even with such unprecedented speed of vaccine and therapeutic development, the pandemic has inflicted devastating worldwide effects, including the loss of millions of human lives, a long-lasting burden on the healthcare system, and economic turmoil. Accelerating actions by several weeks or months can make an enormous difference in an exponentially evolving pandemic, especially at the early stages of infection spread. Therefore, the efficient discovery of effective countermeasures against emerging pathogens can play a critical role in pandemic preparedness for future infectious disease outbreaks.

Antibodies are a major modality for therapy against a wide range of diseases, including infectious diseases, cancer, autoimmunity, cardiovascular, hematologic, neurologic diseases, and others. Antibody discovery efforts have relied on a broad range of technologies, such as hybridoma generation, antigen-specific B cell sorting, phage or yeast display, single-cell B cell sequencing and others^9-14^. Overall, these efforts aim to identify paired heavy–light chain antibody sequences isolated from human or animal samples with prior antigen exposure, followed by monoclonal antibody production and functional characterization. However, with these approaches for antibody discovery, limited information is typically obtained about the antigen specificity of antibody candidates at the screening step. The recent development of the LIBRA-seq (linking B cell receptor to antigen specificity through sequencing) technology enables the simultaneous mapping of antibody sequence to antigen specificity for a large number of antigens in a single next-generation sequencing experiment^15^. LIBRA-seq uses DNA-barcoded antigens to transform B cell receptor (BCR)-antigen binding into a sequence-able event and is of unique utility for discovering broadly reactive antibodies for pathogens that exhibit substantial antigen variability. LIBRA-seq has been applied successfully to identify broadly reactive antibodies against HIV-1, influenza viruses, and coronaviruses^15, 16^.

Although using antigen binding alone as a readout can result in the successful identification of functional antibodies, the discovery process can be inefficient, requiring time-intensive subsequent monoclonal antibody validation steps. This limitation was exemplified by the SARS-CoV-2 antibody discovery initiatives over the past year. Although these efforts have met with notable success, resulting in numerous antibodies and antibody cocktails in clinical trials or approved for emergency use^17-19^, testing of large numbers of antibodies (frequently hundreds to thousands) was generally required. Typically, a small fraction of tested antibodies was neutralizing, with hit rates in the range of 0.3 to 23% in various studies^2-7^.

To overcome this limitation, we developed LIBRA-seq with ligand blocking, a second-generation LIBRA-seq technology that incorporates a functional readout into the antibody discovery process. To achieve this goal, a ligand and its cognate target antigen(s) are each labeled with a unique oligonucleotide barcode (**Supplementary Figure 1A**), enabling the transformation of antigen-ligand interactions into sequence-able events. This allows for the identification of B cells that have high LIBRA-seq scores for the target antigen(s) and low LIBRA-seq scores for the ligand, as B cells that are likely able to potently block antigen-ligand interactions (**Supplementary Figure 1A**). Therefore, a single high-throughput LIBRA-seq with ligand blocking experiment provides both binding (antigen recognition) and functional (ligand blocking) information for thousands of B cells at a time, at the single-cell level.

As a proof of concept, we performed antibody discovery for SARS-CoV-2-specific antibodies from the peripheral B cells from convalescent subjects with past SARS-CoV-2 infection. This provides a suitable opportunity for validating the LIBRA-seq with ligand blocking technology since antibodies that block the interactions of the SARS-CoV-2 spike (S) protein with its receptor angiotensin-converting enzyme 2 (ACE2) are among the most potently neutralizing identified to date^2-7, 17-19^. We performed three separate LIBRA-seq experiments, with screening libraries that included: experiment 1, ACE2 and SARS-CoV-2 S; experiment 2, a titration series of different aliquots of SARS-CoV-2 S, each labeled with a unique barcode; and experiment 3, ACE2 and a titration series of S (**Figure 1A**). The incorporation of a titration series of the S antigen in the LIBRA-seq screening library for experiments 2 and 3 aimed to assess the strength of BCR-antigen interactions through a sequencing readout (**Supplementary Figure 1B, C**). The antigen screening library for each of the experiments also included an influenza virus hemagglutinin and an HIV-1 envelope variant protein as controls.

**Figure 1.**
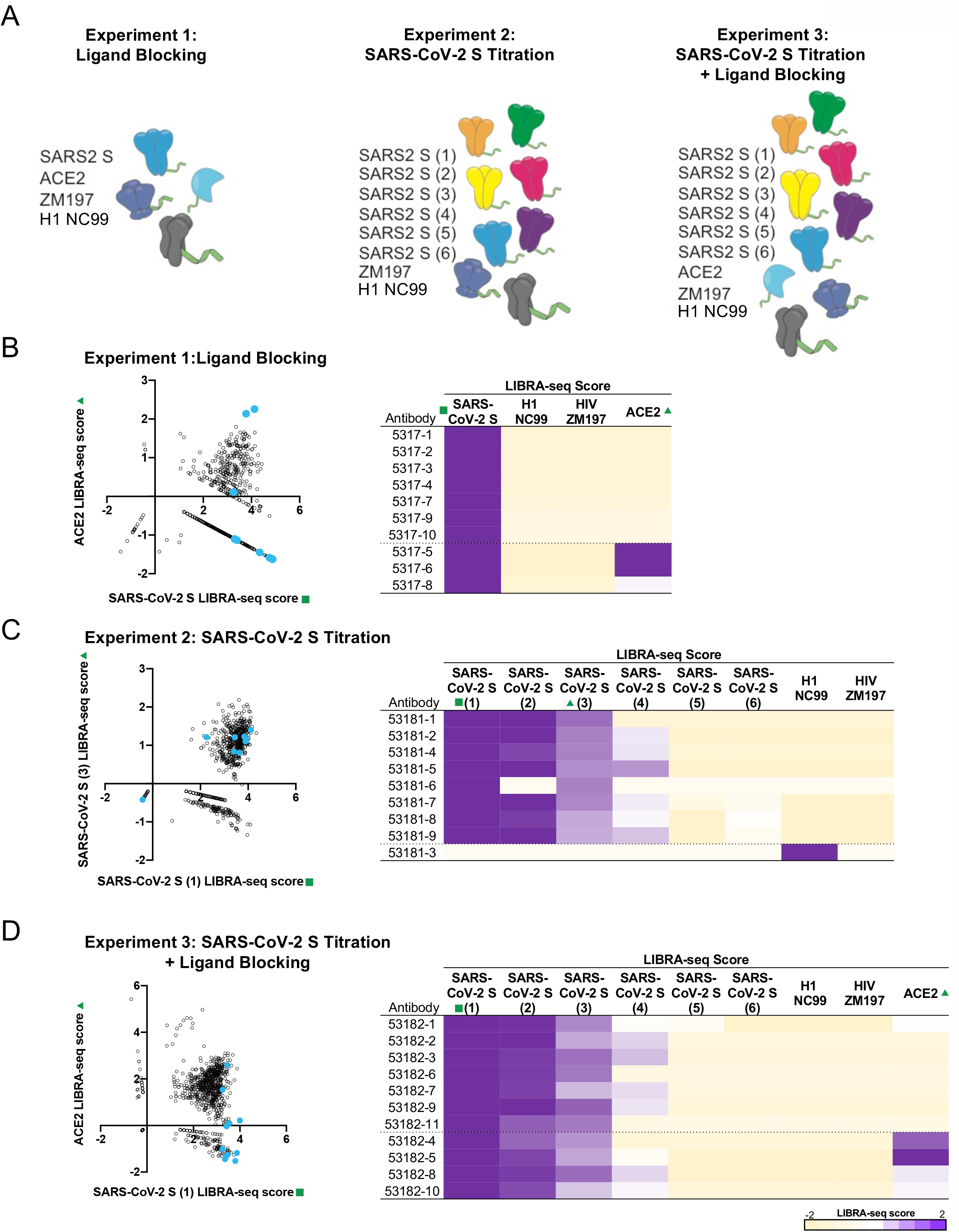
Recovery of antibodies using LIBRA-seq with ligand blocking. **A**. Experimental setup of three LIBRA-seq experiments: experiment 1, LIBRA-seq with ligand blocking; experiment 2, LIBRA-seq with a SARS-CoV-2 S titration; and experiment 3, LIBRA-seq with a SARS-CoV-2 S titration and ligand blocking. After next-generation sequencing, hundreds of B cells were recovered that had paired heavy/light chain sequencing information and antigen reactivity information for the three experiments. For experiment 1 (**B**), 2 (**C**), and 3 (**D**), LIBRA-seq scores for all cells per experiment are shown as open circles (n=828, 829, 957, respectively). Antibodies chosen for expression and validation are highlighted in light blue. LIBRA-seq scores for antigens in each experimental library for antibodies expressed are shown from -2 to 2, as tan to purple, respectively. Scores outside of this range are shown as the minimum and maximum values. For experiment 2 and 3, six different aliquots of S protein were added in a titration series (1-6).

The application of LIBRA-seq resulted in 829, 830, and 957 antigen-specific B cells for the three experiments, respectively. We prioritized a set of B cells for monoclonal antibody production and validation based on the following conditions: for experiments 1 and 3 where ACE2 was included in the screening library, we selected B cells with high LIBRA-seq scores for S and low scores for ACE2; and for experiment 2, we selected B cells that had positive scores for multiple aliquots of S (**Figure 1B-D**). B cells with high S and high ACE2 LIBRA-seq scores were also selected as controls from experiments 1 and 3, along with an influenza-specific B cell from experiment 2 (**Figure 1B-D**). Although some of the selected antibodies appeared to be clonally related, overall they exhibited diverse sequence features, including V_H_- and J_H_-gene usage, HCDR3 lengths, and somatic mutation numbers for both the heavy and light chains (**Supplementary Figure 2A**).

To confirm the predicted antigen specificity for the selected antibodies, we tested them for binding to S ectodomain by ELISA (**Figure 2A, Supplementary Figure 2B**). The LIBRA-seq predicted antigen specificity was confirmed for 26 of 27 (96%) antibodies, including 25 SARS-CoV-2 antibodies and 1 influenza virus antibody. Next, to map the general antibody epitope regions, we also tested the antibodies for binding to S1, S2, receptor-binding domain (RBD), and N-terminal domain (NTD) proteins (**Figure 2A, Supplementary Figure 2B**). The majority of the antibodies from experiments 1 and 3 recognized the RBD, whereas none of the antibodies from experiment 2 recognized the RBD, suggesting that the inclusion of a sequencing readout for ACE2 blocking impacted the ability to select antibodies with target epitope specificities (**Figure 2A, Supplementary Figure 2B**). The antibodies had a wide range of affinities for RBD or NTD, including several ultra-potent antibodies (*K*_D_ <1 nM) from experiments 1 and 3 (**Figure 2B**). Next, we tested the ability of the antibodies to block ACE2 binding to spike. For the antibodies predicted to block ACE2 by LIBRA-seq, 57% from experiment 1 and 67% from experiment 3 demonstrated ACE2 blocking ability via ELISA (**Figure 2C, Supplementary Figure 2C**). In contrast, no antibodies from experiment 2 blocked ACE2 binding (**Figure 2C, Supplementary Figure 2C**). In addition, we tested the antibodies in a VSV SARS-CoV-2 chimeric virus neutralization assay. For the antibodies predicted to block ACE2 by LIBRA-seq, 86% from experiment 1 and 67% from experiment 3 were neutralizing (**Figure 2D-F**). In contrast, only two clonally related antibodies (25%) from experiment 2 were neutralizing (**Figure 2D-F, Supplementary Figure 2D**). These results highlight the importance of including ligand blocking in the LIBRA-seq screening for selectively identifying potently neutralizing antibodies. Significant differences were not detected for the neutralization potencies between antibodies from experiments 1 and 3 (p-value: >0.99, ANOVA with the Kruskal-Wallis Test; **Figure 2E**), suggesting that antigen titrations did not have a significant effect on the ability to identify potently neutralizing antibodies.

**Figure 2.**
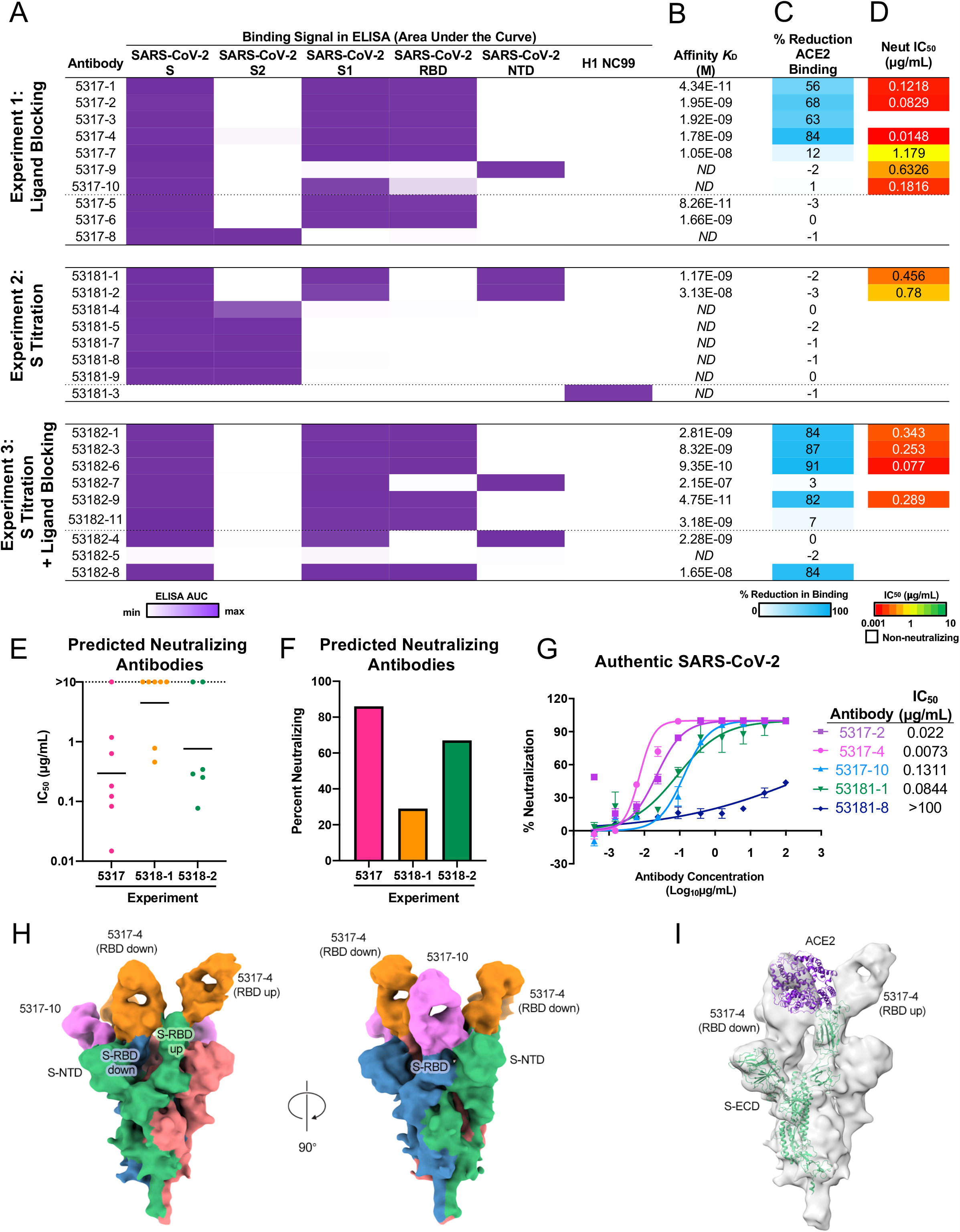
Validation and characterization of expressed antibodies. Names of antibodies that were expressed and their characteristics are shown. Each row represents an antibody. **A**. ELISA area under the curve (AUC) values for binding to SARS-CoV-2 recombinant antigen proteins and a negative control influenza hemagglutinin protein are shown for antibodies in each experiment calculated from data in Supplementary Figure 2B. **B**. Affinity (*K*_D_) of antibodies for SARS-CoV-2 RBD or NTD (based on epitope shown in A) was determined by biolayer interferometry. ND, not done. **C**. Percent reduction in ACE2 binding by ELISA is shown as a heatmap from 0 to 100% reduction in binding as white to blue respectively. **D**. VSV SARS-CoV-2 neutralization IC_50_ values are shown as a heatmap from high potency (red) to low potency (green). Non-neutralizing antibodies are shown as white. For **E** and **F**, “Predicted Neutralizing Antibodies” were defined as the subset of selected antibodies with low ACE2 LIBRA-seq scores from experiments 1 (7 antibodies) and 3 (6 antibodies), and all antibodies with high LIBRA-seq scores for SARS-CoV-2 S from experiment 2 (7 antibodies). **E**. For these predicted neutralizers, plotted are the IC_50_ values (µg/mL) for SARS-CoV-2 neutralization by real time cell analysis (RTCA) with VSV-SARS-CoV-2 (IC_50_ value for each antibody shown as single dot). Line shown is geometric mean. Non-neutralizing antibodies are shown as >10 µg/mL. **F**. The percent of neutralizing antibodies from the set of predicted neutralizers is shown for each experiment. **G**. Authentic SARS-CoV-2 neutralization for a panel of antibodies is shown. Data are represented as mean ± SD. IC_50_ values are shown to the right of the panel. **H**. 10 Å-resolution cryo-EM structure of Fab-spike complex for 5317-4 Fab (orange) and 5317-10 Fab (pink). Spike protomers are shown in green, blue, and red. **I**. Fab-spike complex structure modeled with ACE2 (purple).

Next, we tested a panel of antibodies for neutralization of authentic SARS-CoV-2 virus in a plaque reduction neutralization test and found that, just as in the VSV assay, antibody 5317-4 exhibited the most potent neutralization, with an IC_50_ value of 7.3 ng/mL (**Figure 2G**), on par with antibodies currently in clinical trials or approved for emergency use^17-19^. In addition, to investigate the recognition of SARS-CoV-2 S by LIBRA-seq-identified antibodies, we determined a 9 Å-resolution Cryo-EM structure of the antigen-binding fragments (Fabs) of 5317-4 and another neutralizing antibody, 5317-10, bound to the SARS-CoV-2 S extracellular domain (ECD) (**Figure 2H**). We chose 5317-4 based on its potent neutralization, high affinity for the S RBD, and ACE2 competition. The 3D reconstruction revealed that 5317-4 binds to the RBD in the “up” and “down” conformations, and its epitope partially overlaps the ACE2 binding footprint (**Figure 2H-I**). In the RBD up position, 5317-4 binds near the receptor-binding motif at an angle that appears to moderately clash with RBD-bound ACE2 (**Figure 2I**). Additionally, when bound to the RBD in the down conformation, 5317-4 competes with ACE2 binding to the adjacent, otherwise accessible RBD in the up position (**Figure 2I**). 5317-10 was of interest due to its inconclusive epitope bin, as it bound to S1, but not to the individual RBD or NTD constructs (**Figure 2A**). The map revealed that 5317-10 binds a novel quaternary epitope that bridges an RBD in the down position and the NTD of an adjacent protomer (**Figure 2H**). This mode of recognition may prevent the RBD from transitioning into an ACE2-accessible up position, thereby preventing binding by ACE2.

Together, the results from the three LIBRA-seq experiments showcase the advantages of including ligand blocking as part of the sequencing readout. The experiment without ligand blocking (experiment 2) had a lower hit rate for identifying virus-neutralizing antibodies. Through a single high-throughput sequencing experiment, LIBRA-seq with ligand blocking identified ultra-potent SARS-CoV-2 antibodies, requiring the subsequent production and validation of less than a dozen antibodies per experiment. The observed hit rates for the discovery of potently neutralizing antibodies are a substantial improvement over what has been reported in the literature, which typically required the screening of hundreds to thousands of antibody candidates to identify a small number of antibodies with similar levels of neutralization potencies to the antibodies described in this manuscript^2-7^. The ability to save weeks or months during the antibody lead discovery process can have an extraordinary impact on the evolution of an infectious disease outbreak, especially in the early stages of infection spread. As such, the application of LIBRA-seq with ligand blocking for the efficient discovery of highly potent antibodies can provide critical advantages for rapid development of therapeutic and preventive countermeasures. More broadly, LIBRA-seq with ligand blocking presents a general platform for discovering functional antibodies, with applications to virtually any area where targeting the disruption of antigen-ligand interaction is a prime therapeutic goal.

## Acknowledgements

We thank Angela Jones, Latha Raju, and Jamie Roberson of Vanderbilt Technologies for Advanced Genomics for their expertise regarding NGS and library preparation; David Flaherty and Brittany Matlock of the Vanderbilt Flow Cytometry Shared Resource for help with flow panel optimization; and members of the Georgiev laboratory for comments on the manuscript. The Vanderbilt VANTAGE Core provided technical assistance for this work. VANTAGE is supported in part by CTSA grant 5UL1 RR024975-03, the Vanderbilt Ingram Cancer Center (P30 CA68485), the Vanderbilt Vision Center (P30 EY08126), and NIH/NCRR (G20 RR030956). This work was conducted in part using the resources of the Advanced Computing Center for Research and Education at Vanderbilt University (Nashville, TN). Flow cytometry experiments were performed in the VUMC Flow Cytometry Shared Resource. The VUMC Flow Cytometry Shared Resource is supported by the Vanderbilt Ingram Cancer Center (P30 CA68485) and the Vanderbilt Digestive Disease Research Center (DK058404).

For work described in this manuscript, I.S.G., A.R.S., K.J.K., K.A.P., N.R., and L.W. were supported in part by NIH NIAID award R01AI131722-S1, the Hays Foundation COVID-19 Research Fund, Fast Grants, and CTSA award No. *UL1 TR002243* from the National Center for Advancing Translational Sciences. J.S.M, N.J., and D.W. were supported in part by a National Institutes of Health (NIH)/National Institute of Allergy and Infectious Diseases (NIAID) grant awarded to J.S.M. (R01-AI127521), Welch Foundation grant F-0003-19620604, and NIH NIAID award R01AI131722-S1. This work was supported by Defense Advanced Research Projects Agency (DARPA) grant HR0011-18-2-0001, U.S. N.I.H. contract 75N93019C00074, N.I.H. grant R01 AI157155, the Dolly Parton COVID-19 Research Fund at Vanderbilt, and a grant from Fast Grants, Mercatus Center, George Mason University. J.E.C. is a recipient of the 2019 Future Insight Prize from Merck KGaA, which supported this work with a grant.

## Author Contributions

Methodology, A.R.S., I.S., and I.S.G.; Investigation, A.R.S., N.J., K.J.K., N.S., D.W., S.P., K.A.P., N.J., R.N., R.E.S., L.W., J.E.C., A.B., R.H.C., J.S.M., and I.S.G.; Software, A.R.S., N.R.; Validation, A.R.S., K.J.K.; Writing - Original Draft, A.R.S. and I.S.G.; Writing -Review & Editing, all authors; Funding Acquisition, I.S.G., J.E.C., A.B., R.H.C., J.S.M., and A.R.S and K.J.K.; Resources, I.S.G., J.E.C., A.B., R.H.C., and J.S.M..; Supervision, I.S.G.

## Declaration of Interest

A.R.S. and I.S.G. are co-founders of AbSeek Bio. I.S.G., A.R.S, and K.J.K. are listed as inventors on antibodies described herein. I.S.G., A.R.S, and I.S. are listed as inventors on patent applications for the LIBRA-seq technology. J.E.C. has served as a consultant for Luna Biologics, is a member of the Scientific Advisory Board Meissa Vaccines and is Founder of IDBiologics. The Crowe laboratory has received funding support in sponsored research agreements from AstraZeneca, IDBiologics, and Takeda. The Georgiev laboratory at Vanderbilt University Medical Center has received unrelated funding from Takeda Pharmaceuticals.

## Data Availability Statement

All unique reagents generated in this study are available from the corresponding author with a completed Materials Transfer Agreement. Sequences for antibodies identified and characterized in this study will be deposited to GenBank and raw sequencing data will be deposited to Sequence Read Archive and will be available upon publication. Further information and requests for resources and reagents should be directed to the corresponding author, Ivelin Georgiev.

## Code Availability

Custom scripts used to analyze data in this manuscript are available upon request to the corresponding author.

## METHODS

### Donor Information

PBMC samples were purchased from Cellero. The PBMCs were from subjects with past SARS-CoV-2 infection at least 14 days post symptom cessation. For experiment 1, three samples were pooled from donors 523, 527, and 528. For experiments 2 and 3, samples from donor 523 were used for LIBRA-seq. Donor 523 had a plaque reduction neutralization test titer of 1:2,560.

### Antigen Purification

A variety of recombinant soluble protein antigens were used in the LIBRA-seq experiment and other experimental assays.

Plasmids encoding residues 1–1208 of the SARS-CoV-2 spike with a mutated S1/S2 cleavage site, proline substitutions at positions 817, 892, 899, 942, 986 and 987, and a C-terminal T4-fibritin trimerization motif, an 8x HisTag, and a TwinStrepTag (SARS-CoV-2 spike HP); 1−615 of human ACE2 with a C-terminal HRV3C protease cleavage site, a TwinStrepTag and an 8XhisTag (ACE2) were transiently transfected in Expi293F cells using polyethylenimine. Transfected supernatants were harvested 5 days after expression and purified over a StrepTrap column (Cytiva Life Sciences), as previously described. Both recombinant SARS-CoV-2 S HP and ACE2 were further purified to homogeneity using a Superose6 Increase column (Cytiva Life Sciences).

For the HIV-1 gp140 SOSIP variant from strain ZM197 (clade C) and hemagglutinin from strain A/New Caledonia/20/99 (H1N1) (GenBank ACF41878), recombinant, soluble antigens contained an AviTag and were expressed in Expi293F cells using polyethylenimine transfection reagent and cultured. FreeStyle F17 expression medium supplemented with pluronic acid and glutamine was used. The cells were cultured at 37°C with 8% CO_2_ saturation and shaking. After 5-7 days, cultures were centrifuged and supernatant was filtered and run over an affinity column of agarose-bound *Galanthus nivalis* lectin. The column was washed with PBS and antigens were eluted with 30 mL of 1M methyl-a-D-mannopyranoside. Protein elutions were buffer exchanged into PBS, concentrated, and run on a Superdex 200 Increase 10/300 GL Sizing column on the AKTA FPLC system. Fractions corresponding to correctly folded protein were collected, analyzed by SDS-PAGE and antigenicity was characterized by ELISA using known monoclonal antibodies specific to each antigen. AviTagged antigens were biotinylated using BirA biotin ligase (Avidity LLC).

SARS-CoV-2 S1, SARS-CoV-2 S2, SARS-CoV-2 RBD and SARS-CoV-2 NTD proteins were purchased from the commercial vendor, Sino Biological.

### DNA-barcoding of Antigens

We used oligos that possess 15 bp antigen barcode, a sequence capable of annealing to the template switch oligo that is part of the 10X bead-delivered oligos and contain truncated TruSeq small RNA read 1 sequences in the following structure: 5′-CCTTGGCACCCGAGAATTCCANNNNNNNNNNNNNCCCATATAAGA*A*A-3′, where Ns represent the antigen barcode as previously described^15^. For each antigen, a unique DNA barcode was directly conjugated to the antigen itself. In particular, 5′ amino-oligonucleotides were conjugated directly to each antigen using the SoluLINK Protein-Oligonucleotide Conjugation Kit (TriLink cat no. S-9011) according to manufacturer’s instructions. Briefly, the oligo and protein were desalted, and then the amino-oligo was modified with the 4FB crosslinker, and the biotinylated antigen protein was modified with S-HyNic. Then, the 4FB-oligo and the HyNic-antigen were mixed. This process causes a stable bond to form between the protein and the oligonucleotide. The concentration of the antigen-oligo conjugates was determined by a BCA assay, and the HyNic molar substitution ratio of the antigen-oligo conjugates was analyzed using the NanoDrop according to the SoluLINK protocol guidelines. AKTA FPLC was used to remove excess oligonucleotide from the protein-oligo conjugates, which were also verified using SDS-PAGE with a silver stain. Antigen-oligo conjugates were also used in flow cytometric titration experiments.

### Antigen-specific B Cell Sorting

Cells were stained and mixed with DNA-barcoded antigens and other antibodies, and then sorted using fluorescence activated cell sorting (FACS). First, cells were counted, and viability was assessed using trypan blue. Then, cells were washed three times with DPBS supplemented with 0.1% bovine serum albumin (BSA). Cells were resuspended in DPBS-BSA and stained with cell markers including viability dye (Ghost Red 780), CD14-APC-Cy7, CD3-FITC, CD19-BV711, and IgG-PE-Cy5. Additionally, antigen-oligo conjugates were added to the stain. After staining in the dark for 30 minutes at room temperature, cells were washed three times with DPBS-BSA at 300 x g for five minutes. Cells were then incubated for 15 minutes at room temperature with Streptavidin-PE to label cells with bound antigen. Cells were washed three times with DPBS-BSA, resuspended in DPBS, and sorted by FACS. Antigen positive cells were bulk sorted and delivered to the Vanderbilt Technologies for Advanced Genomics (VANTAGE) sequencing core at an appropriate target concentration for 10X Genomics library preparation and subsequent sequencing. Flow cytometry data were analyzed using FlowJo.

### Sample Preparation, Library Preparation, and Sequencing

Single-cell suspensions were loaded onto the Chromium Controller microfluidics device (10X Genomics) and processed using the B-cell Single Cell V(D)J solution according to manufacturer’s suggestions for a target capture of 10,000-20,000 B cells, with minor modifications to intercept, amplify and purify the antigen barcode libraries as previously described^15^.

### Sequence processing and bioinformatic analysis

We used our previously described pipeline to use paired-end FASTQ files of oligo libraries as input, process and annotate reads for cell barcode, UMI, and antigen barcode, and generate a cell barcode - antigen barcode UMI count matrix^15, 16^. BCR contigs were processed using Cell Ranger (10X Genomics) using GRCh38 as reference. Antigen barcode libraries were also processed using Cell Ranger (10X Genomics). The overlapping cell barcodes between the two libraries were used as the basis of the subsequent analysis. We removed cell barcodes that had only non-functional heavy chain sequences as well as cells with multiple functional heavy chain sequences and/or multiple functional light chain sequences, reasoning that these may be multiplets. Additionally, we aligned the BCR contigs (filtered_contigs.fasta file output by Cell Ranger, 10X Genomics) to IMGT reference genes using HighV-Quest^20^. The output of HighV-Quest was parsed using ChangeO^21^ and merged with an antigen barcode UMI count matrix. Finally, we determined the LIBRA-seq score for each antigen in the library by calculating the centered-log ratios (CLR) of each antigen UMI count for each cell as previously described^15^.

### Antibody Expression and Purification

For each antibody, variable genes were inserted into custom plasmids encoding the constant region for the IgG1 heavy chain as well as respective lambda and kappa light chains (pTwist CMV BetaGlobin WPRE Neo vector, Twist Bioscience). Antibodies were expressed in Expi293F mammalian cells (Thermo Fisher Scientific) by co-transfecting heavy chain and light chain expressing plasmids using polyethylenimine transfection reagent and cultured for 5 to 7 days. Cells were maintained in FreeStyle F17 expression medium supplemented at final concentrations of 0.1% Pluronic Acid F-68 and 20% 4 mM L-Glutamine. These cells were cultured at 37°C with 8% CO_2_ saturation and shaking. After transfection and 5-7 days of culture, cell cultures were centrifuged and supernatant was 0.45 μm filtered with Nalgene Rapid Flow Disposable Filter Units with PES membrane. Filtered supernatant was run over a column containing Protein A agarose resin equilibrated with PBS. The column was washed with PBS, and then antibodies were eluted with 100 mM Glycine HCl at 2.7 pH directly into a 1:10 volume of 1M Tris-HCl pH 8.0. Eluted antibodies were buffer exchanged into PBS 3 times using Amicon Ultra-centrifugal filter units and concentrated. Antibody plasmids were sequenced. If antibody sequences did not match expected heavy or light chain, antibody was excluded from downstream analysis.

### High-throughput Antibody Expression

For high-throughput production of recombinant antibodies, approaches were used that are designated as microscale. For antibody expression, microscale transfection was performed (∼1 mL per antibody) of CHO cell cultures using the Gibco ExpiCHO Expression System and a protocol for deep 96-well blocks (Thermo Fisher Scientific). In brief, synthesized antibody-encoding DNA (∼2 μg per transfection) was added to OptiPro serum free medium (OptiPro SFM), incubated with ExpiFectamine CHO Reagent and added to 800 µL of ExpiCHO cell cultures into 96-deep-well blocks using a ViaFlo 384 liquid handler (Integra Biosciences). The plates were incubated on an orbital shaker at 1,000 r.p.m. with an orbital diameter of 3 mm at 37°C in 8% CO_2_. The next day after transfection, ExpiFectamine CHO Enhancer and ExpiCHO Feed reagents (Thermo Fisher Scientific) were added to the cells, followed by 4 d incubation for a total of 5 d at 37°C in 8% CO_2_. Culture supernatants were collected after centrifuging the blocks at 450 x *g* for 5 min and were stored at 4°C until use. For high-throughput microscale antibody purification, fritted deep-well plates were used containing 25 μL of settled protein G resin (GE Healthcare Life Sciences) per well. Clarified culture supernatants were incubated with protein G resin for antibody capturing, washed with PBS using a 96-well plate manifold base (Qiagen) connected to the vacuum and eluted into 96-well PCR plates using 86 μL of 0.1 M glycine-HCL buffer pH 2.7. After neutralization with 14 μL of 1 M Tris-HCl pH 8.0, purified antibodies were buffer-exchanged into PBS using Zeba Spin Desalting Plates (Thermo Fisher Scientific) and stored at 4°C until use.

### ELISA

To assess antibody binding, soluble protein was plated at 2 μg/mL overnight at 4°C. The next day, plates were washed three times with PBS supplemented with 0.05% Tween-20 (PBS-T) and coated with 5% milk powder in PBS-T. Plates were incubated for one hour at room temperature and then washed three times with PBS-T. Primary antibodies were diluted in 1% milk in PBS-T, starting at 10 μg/mL with a serial 1:5 dilution and then added to the plate. The plates were incubated at room temperature for one hour and then washed three times in PBS-T. The secondary antibody, goat anti-human IgG conjugated to peroxidase, was added at 1:10,000 dilution in 1% milk in PBS-T to the plates, which were incubated for one hour at room temperature. Plates were washed three times with PBS-T and then developed by adding TMB substrate to each well. The plates were incubated at room temperature for ten minutes, and then 1N sulfuric acid was added to stop the reaction. Plates were read at 450 nm.

Data are represented as mean ± SEM for one ELISA experiment. ELISAs were repeated 2 or more times. If ELISA replicates were inconsistent over more than three experiments, antibody was excluded from in vitro characterization analysis. The area under the curve (AUC) was calculated using Prism software version 8.0.0 (GraphPad).

### ACE2 Binding Inhibition Assay

96-well plates were coated with 2 μg/mL purified recombinant SARS-CoV-2 at 4°C overnight. The next day, plates were washed three times with PBS supplemented with 0.05% Tween-20 (PBS-T) and coated with 5% milk powder in PBS-T. Plates were incubated for one hour at room temperature and then washed three times with PBS-T. Purified anti were diluted in blocking buffer at 10 μg/mL in triplicate, added to the wells, and incubated at room temperature. Without washing, recombinant human ACE2 protein with a mouse Fc tag was added to wells for a final 0.4 μg/mL concentration of ACE2 and incubated for 40 minutes at room temperature. Plates were washed three times with PBS-T, and bound ACE2 was detected using HRP-conjugated anti-mouse Fc antibody and TMB substrate. The plates were incubated at room temperature for ten minutes, and then 1N sulfuric acid was added to stop the reaction. Plates were read at 450 nm. ACE2 binding without antibody served as a control. Experiment was done in biological replicate and technical triplicates.

### BioLayer Interferometry (BLI)

Purified antibodies were immobilized to AHC sensortips (FortéBio) to a response level of approximately 1.4 nm in a buffer composed of 10 mM HEPES pH 7.5, 150 mM NaCl, 3 mM EDTA, 0.05% Tween 20 and 0.1% (w/v) BSA. Immobilized antibodies were then dipped into wells containing two-fold dilutions of either SARS-CoV-2 RBD-SD1 (residues 306–577) or SARS-CoV-2 NTD, ranging in concentration from 10–0.156 nM, to measure association kinetics. Dissociation kinetics were measured by dipping sensortips into wells containing only buffer. Data were reference subtracted and kinetics were calculated in Octet Data Analysis software v10.0 using a 1:1 binding model.

### RTCA Method for Initial Screening of Antibody Neutralizing Activity

To screen for neutralizing activity in the panel of recombinantly expressed antibodies, we used a high-throughput and quantitative RTCA assay and xCelligence RTCA HT Analyzer (ACEA Biosciences) that assesses kinetic changes in cell physiology, including virus-induced cytopathic effect (CPE). Twenty µL of cell culture medium (DMEM supplemented with 2% FBS) was added to each well of a 384-well E-plate using a ViaFlo384 liquid handler (Integra Biosciences) to obtain background reading. Six thousand (6,000) Vero-furin cells in 20 μL of cell culture medium were seeded per well, and the plate was placed on the analyzer. Sensograms were visualized using RTCA HT software version 1.0.1 (ACEA Biosciences). For a screening neutralization assay, equal amounts of virus were mixed with micro-scale purified antibodies in a total volume of 40 μL using DMEM supplemented with 2% FBS as a diluent and incubated for 1 h at 37°C in 5% CO_2_. At ∼17–20 h after seeding the cells, the virus–antibody mixtures were added to the cells in 384-well E-plates. Wells containing virus only (in the absence of antibody) and wells containing only Vero cells in medium were included as controls. Plates were measured every 8–12 h for 48–72 h to assess virus neutralization. Micro-scale antibodies were assessed in four 5-fold dilutions (starting from a 1:20 sample dilution), and their concentrations were not normalized. Neutralization was calculated as the percent of maximal cell index in control wells without virus minus cell index in control (virus-only) wells that exhibited maximal CPE at 40–48 h after applying virus–antibody mixture to the cells. An antibody was classified as fully neutralizing if it completely inhibited SARS-CoV-2-induced CPE at the highest tested concentration, while an antibody was classified as partially neutralizing if it delayed but did not fully prevent CPE at the highest tested concentration^2, 22^.

### Real-time Cell Analysis (RTCA) Neutralization Assay

To determine neutralizing activity of IgG, we used real-time cell analysis (RTCA) assay on an xCELLigence RTCA MP Analyzer (ACEA Biosciences Inc.) that measures virus-induced cytopathic effect (CPE)^23^. Briefly, 50 μL of cell culture medium (DMEM supplemented with 2% FBS) was added to each well of a 96-well E-plate using a ViaFlo384 liquid handler (Integra Biosciences) to obtain background reading. A suspension of 18,000 Vero-E6 cells in 50 μL of cell culture medium was seeded in each well, and the plate was placed on the analyzer. Measurements were taken automatically every 15 min, and the sensograms were visualized using RTCA software version 2.1.0 (ACEA Biosciences Inc). VSV-SARS-CoV-2 (0.01 MOI, ∼120 PFU per well) was mixed 1:1 with a dilution of antibody in a total volume of 100 μL using DMEM supplemented with 2% FBS as a diluent and incubated for 1 h at 37°C in 5% CO_2_. At 16 h after seeding the cells, the virus-antibody mixtures were added in replicates to the cells in 96-well E-plates. Triplicate wells containing virus only (maximal CPE in the absence of antibody) and wells containing only Vero cells in medium (no-CPE wells) were included as controls. Plates were measured continuously (every 15 min) for 48 h to assess virus neutralization. Normalized cellular index (CI) values at the endpoint (48 h after incubation with the virus) were determined using the RTCA software version 2.1.0 (ACEA Biosciences Inc.). Results are expressed as percent neutralization in a presence of respective antibody relative to control wells with no CPE minus CI values from control wells with maximum CPE. RTCA IC_50_ values were determined by nonlinear regression analysis using Prism software.

### Plaque Reduction Neutralization Test (PRNT)

The virus neutralization with live authentic SARS-CoV-2 virus (USA-WA1) was performed in the BSL-3 facility of the Galveston National Laboratory using Vero E6 cells (ATCC CRL-1586) following the standard procedure. Vero E6 cells were cultured in 96-well plates (10^4^ cells/well). Next day, 4-fold serial dilutions of antibodies were made using MEM-2% FBS, as to get an initial concentration of 100 µg/mL. Equal volume of diluted antibodies (60 µL) were mixed gently with original SARS-CoV-2 (USA-WA1) (60 µL containing 200 pfu) and incubated for 1 h at 37°C/5% CO_2_ atmosphere. The virus-serum mixture (100 µL) was added to cell monolayer in duplicates and incubated for 1 at h 37°C/5% CO_2_ atmosphere. Later, the virus-serum mixture was discarded gently, and cell monolayer was overlaid with 0.6% methylcellulose and incubated for 2 days. The overlay was removed, and the plates were fixed in 4% paraformaldehyde twice following BSL-3 protocol. The plates were stained with 1% crystal violet and virus-induced plaques were counted. The percent neutralization and/or NT_50_ of antibody was calculated by dividing the plaques counted at each dilution with plaques of virus-only control. For antibodies, the inhibitory concentration at 50% (IC_50_) values were calculated in Prism software (GraphPad) by plotting the midway point between the upper and lower plateaus of the neutralization curve among dilutions.

### Fab Preparation

To generate Fabs, IgGs were incubated with Lys-C at 1:4,000 (weight:weight) overnight at 37 °C. EDTA free protease inhibitor (Roche) was dissolved to 25X and then added to the sample at a final 1X concentration. The sample was passed over a Protein A column. The flow-through was collected run on a Superdex 200 Increase 10/300 GL Sizing column on the AKTA FPLC system. Fabs were visualized on SDS-PAGE.

### Electron Microscopy Sample Preparation and Data Collection

Purified SARS-CoV-2 S HexaPro ectodomain^24^ and Fabs 5317-4 and 5317-10 were combined at a final complex concentration of 0.4 mg/mL. Fab 5317-10 was added to spike and incubated on ice for 30 minutes before the addition of Fab 5317-4 immediately prior to grid deposition and freezing. The complex was deposited on Au-300 1.2/1.3 grids that had been plasma cleaned for 4 minutes in a Solarus 950 plasma cleaner (Gatan) with a 4:1 ratio of O_2_/H_2_. Excess liquid was blotted for 3 seconds with a force of -4 using a Vitrobot Mark IV (Thermo Fisher) and plunge frozen into liquid ethane. 2,655 micrographs were collected from a single grid with the stage at a 30° tilt using a Titan Krios (Thermo Fisher) equipped with a K3 detector (Gatan). Movies were collected using SerialEM^25^ at 29,000X magnification with a corresponding calibrated pixel size of 0.81 Å/ pixel.

### Cryogenic Electron Microscopy (Cryo-EM)

Motion correction, CTF estimation, particle picking, and 2D classification were performed using cryoSPARC v3.2.0^26^. The final iteration of 2D class averaging distributed 17,710 particles into 50 classes using an uncertainty factor of 3. From that, 13,232 particles were selected and an ab inito reconstruction was performed with four classes followed by heterogeneous refinement of those four classes. 6,803 particles from the highest-quality class were used for homogenous refinement of the best volume without imposed symmetry. The resulting volume was used for an additional round of homogenous refinement. To filter out additional junk particles, an *ab initio* reconstruction was performed with three classes followed by heterogeneous refinement of those three classes. 5,171 particles from the highest-quality class were used for homogenous refinement of the best volume without imposed symmetry, resulting in a final 9 Å map.

### QUANTIFICATION AND STATISTICAL ANALYSIS

ELISA error bars (standard error of the mean) were calculated using GraphPad Prism version 8.0.0. ANOVA analysis was performed for neutralization potency comparisons using GraphPad Prism version 8.0.0.

## FIGURE CAPTIONS

**Supplementary Figure 1.**
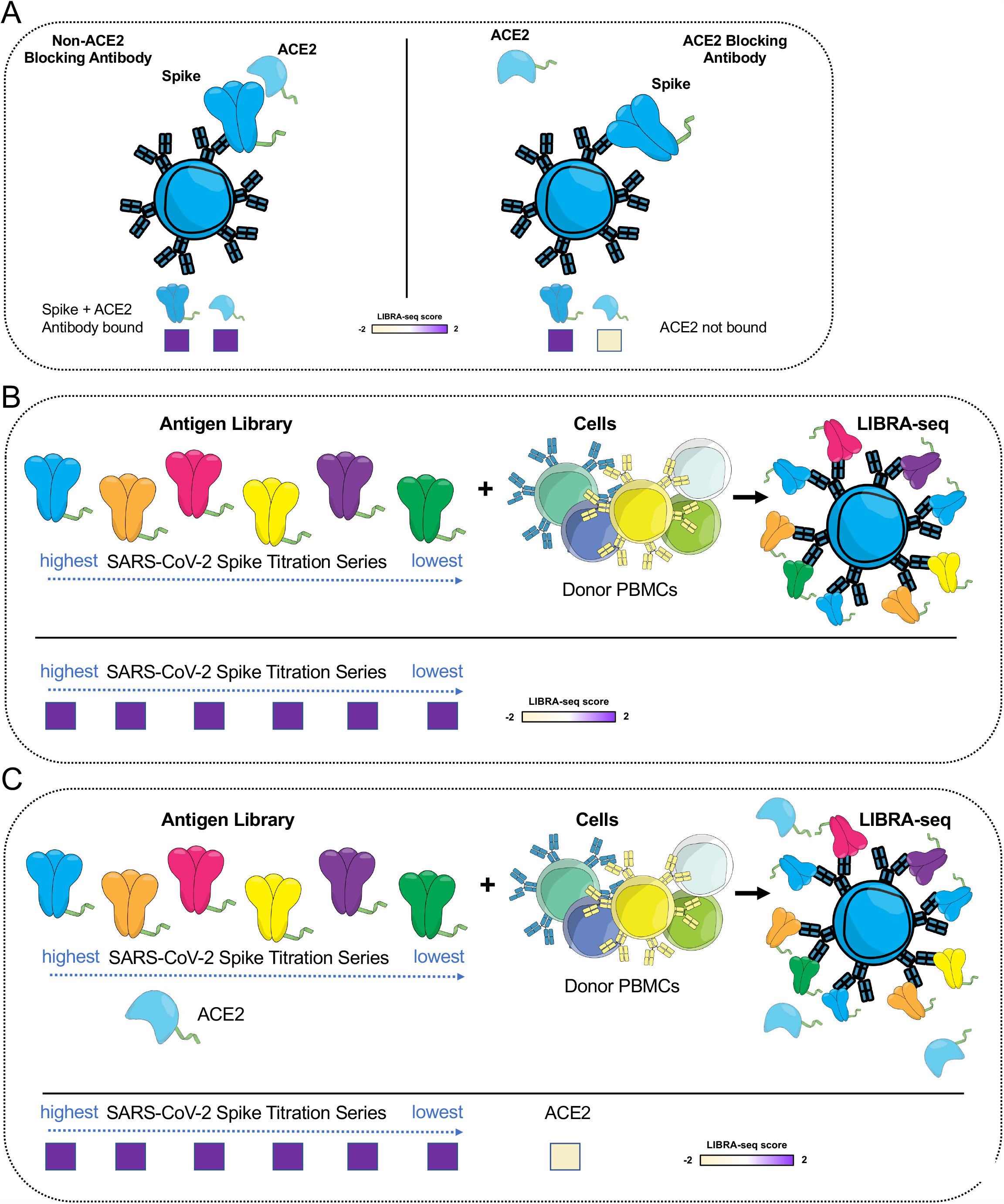
Set up of LIBRA-seq experiments. **A**. An antigen screening library of oligonucleotide-labeled antigens was generated. This library consisted of SARS-CoV-2 spike antigens and negative controls. Additionally, oligo-labeled ACE2 (the SARS-CoV-2 spike host cell receptor) was included. The antigen screening library was mixed with donor PBMCs. This approach allowed for assessment of B cell ligand blocking functionality from the sequencing experiment. **B**. We applied an antigen screening library containing an antigen titration with a goal of identifying high affinity antibodies from LIBRA-seq. In this experiment, six different amounts of oligo-labeled SARS-CoV-2 S protein, each labeled with a different barcode, were included in a screening library, with the hypothesis that antibodies with high affinity for SARS-CoV-2 S would show reactivity (high LIBRA-seq score) even with the S protein added in lower amounts. **C**. Schematic of LIBRA-seq with S titrations and ACE2 included for ligand blocking.

**Supplementary Figure 2.**
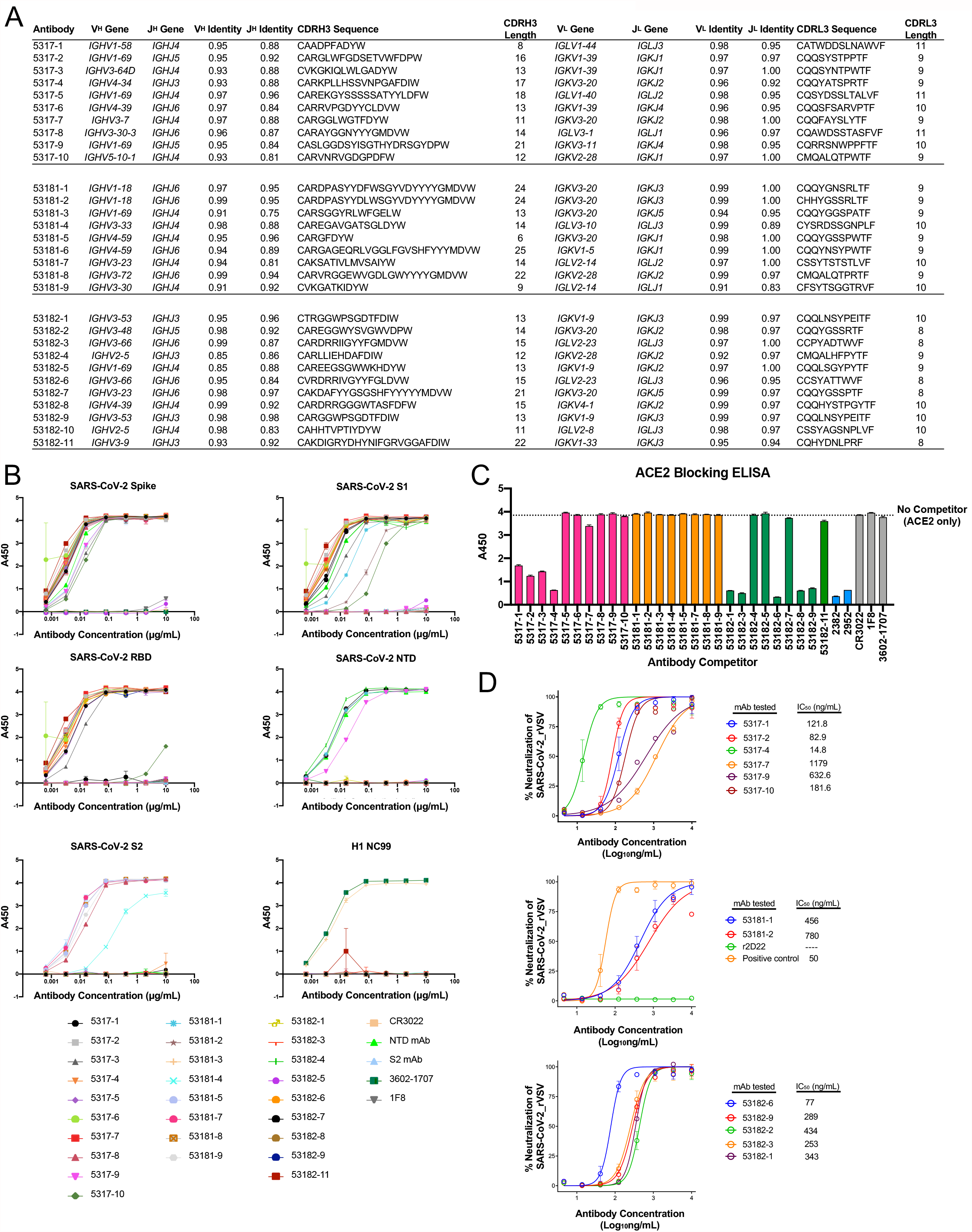
Characterization of LIBRA-seq identified antibodies. **A**. Genetic characteristics for monoclonal antibodies prioritized for expression and validation. V_H_, J_H_, V_L_, J_L_ inferred gene segment identity is shown at the nucleotide level. CDRH3 and CDRL3 amino acid sequence and length are shown. **B**. ELISA binding of antibodies to SARS-CoV-2 spike, SARS-CoV spike, SARS-CoV-2 RBD, SARS-CoV-2 NTD, SARS-CoV-2 S2 and influenza virus H1 NC99. ELISAs were performed in technical duplicates and repeated at least two times. Data are represented as mean ± SEM. **C**. ACE2 blocking ELISA. Antibodies were added to spike and recombinant ACE2 was added and detected. Antibodies that block ACE2 binding show a reduction in absorbance compared to ACE2 binding without competitor (dotted line). ELISAs were performed at one antibody concentration in triplicate and were repeated at least twice. Data are represented as mean ± SEM. **D**. Antibodies were tested in a VSV SARS-CoV-2 RTCA neutralization assay. Neutralization curves and IC_50_ values are shown. Data are represented as mean ± SD.

## References

1. Jiang, S., Hillyer, C. & Du, L. Neutralizing Antibodies against SARS-CoV-2 and Other Human Coronaviruses. Trends Immunol 41, 355–359 (2020).

2. Zost, S.J. et al. Potently neutralizing and protective human antibodies against SARS-CoV-2. Nature 584, 443–449 (2020).

3. Chi, X. et al. A neutralizing human antibody binds to the N-terminal domain of the Spike protein of SARS-CoV-2. Science 369, 650–655 (2020).

4. Wec, A.Z. et al. Broad neutralization of SARS-related viruses by human monoclonal antibodies. Science 369, 731–736 (2020).

5. Brouwer, P.J.M. et al. Potent neutralizing antibodies from COVID-19 patients define multiple targets of vulnerability. Science 369, 643–650 (2020).

6. Rogers, T.F. et al. Isolation of potent SARS-CoV-2 neutralizing antibodies and protection from disease in a small animal model. Science 369, 956–963 (2020).

7. Hansen, J. et al. Studies in humanized mice and convalescent humans yield a SARS-CoV-2 antibody cocktail. Science 369, 1010–1014 (2020).

8. Krammer, F. SARS-CoV-2 vaccines in development. Nature 586, 516–527 (2020).

9. Crowe, J.E., Jr. Principles of Broad and Potent Antiviral Human Antibodies: Insights for Vaccine Design. Cell Host Microbe 22, 193–206 (2017).

10. Scheid, J.F. et al. Broad diversity of neutralizing antibodies isolated from memory B cells in HIV-infected individuals. Nature 458, 636–640 (2009).

11. Wu, X. et al. Rational design of envelope identifies broadly neutralizing human monoclonal antibodies to HIV-1. Science 329, 856–861 (2010).

12. Busse, C.E., Czogiel, I., Braun, P., Arndt, P.F. & Wardemann, H. Single-cell based high-throughput sequencing of full-length immunoglobulin heavy and light chain genes. Eur J Immunol 44, 597–603 (2014).

13. DeKosky, B.J. et al. High-throughput sequencing of the paired human immunoglobulin heavy and light chain repertoire. Nat Biotechnol 31, 166–169 (2013).

14. Tan, Y.C. et al. Barcode-enabled sequencing of plasmablast antibody repertoires in rheumatoid arthritis. Arthritis Rheumatol 66, 2706–2715 (2014).

15. Setliff, I. et al. High-Throughput Mapping of B Cell Receptor Sequences to Antigen Specificity. Cell 179, 1636–1646 e1615 (2019).

16. Shiakolas, A.R. et al. Cross-reactive coronavirus antibodies with diverse epitope specificities and Fc effector functions. Cell Rep Med, 100313 (2021).

17. Chen, P. et al. SARS-CoV-2 Neutralizing Antibody LY-CoV555 in Outpatients with Covid-19. N Engl J Med 384, 229–237 (2021).

18. Cohen, M.S. Monoclonal Antibodies to Disrupt Progression of Early Covid-19 Infection. N Engl J Med 384, 289–291 (2021).

19. Weinreich, D.M. et al. REGN-COV2, a Neutralizing Antibody Cocktail, in Outpatients with Covid-19. N Engl J Med 384, 238–251 (2021).

20. Alamyar, E., Duroux, P., Lefranc, M.P. & Giudicelli, V. IMGT((R)) tools for the nucleotide analysis of immunoglobulin (IG) and T cell receptor (TR) V-(D)-J repertoires, polymorphisms, and IG mutations: IMGT/V-QUEST and IMGT/HighV-QUEST for NGS. Methods Mol Biol 882, 569–604 (2012).

21. Gupta, N.T. et al. Change-O: a toolkit for analyzing large-scale B cell immunoglobulin repertoire sequencing data. Bioinformatics 31, 3356–3358 (2015).

22. Gilchuk, P. et al. Integrated pipeline for the accelerated discovery of antiviral antibody therapeutics. Nat Biomed Eng 4, 1030–1043 (2020).

23. Suryadevara, N. et al. Neutralizing and protective human monoclonal antibodies recognizing the N-terminal domain of the SARS-CoV-2 spike protein. Cell 184, 2316–2331 e2315 (2021).

24. Hsieh, C.L. et al. Structure-based design of prefusion-stabilized SARS-CoV-2 spikes. Science 369, 1501–1505 (2020).

25. Mastronarde, D.N. Automated electron microscope tomography using robust prediction of specimen movements. J Struct Biol 152, 36–51 (2005).

26. Punjani, A., Rubinstein, J.L., Fleet, D.J. & Brubaker, M.A. cryoSPARC: algorithms for rapid unsupervised cryo-EM structure determination. Nat Methods 14, 290–296 (2017).

